# Improving Accuracy of Nuclei Segmentation by Reducing Histological Image Variability BMI 260 - Spring 2017

**DOI:** 10.1101/296806

**Authors:** Yusuf Roohani, Eric Kiss

## Abstract

Cancer is the second leading cause of death in United States. Early diagnosis of this disease is essential for many types of treatment. Cancer is most accurately observed by pathologists using tissue biopsy. In the past, evaluation of tissue samples was done manually, but to improve efficiency and ensure consistent quality, there has been a push to evaluate these algorithmically. One important task in histological analysis is the segmentation and evaluation of nuclei. Nuclear morphology is important to understand the grade and progression of cancer. Convolutional neural networks (CNN) were used to segment train models for nuclei segmentation. Stains are used to highlight cellular features. However, there is significant variability in imaging of stained slides due to differences in stain, slide preparation and slide storage. This make automated methods challenging to implement across different datasets. This paper evaluates four stain normalization methods to reduce the variability between slides. Nuclear segmentation accuracy was evaluated for each normalized method. Baseline segmentation accuracy was improved by more than 50% of its base value as measured by the AUC and Recall. We believe this is the first study to look at the impact of four stain normalization approaches (histogram equalization, Reinhart, Macenko, Khan) on segmentation accuracy.

## 1 Introduction

Diagnoses made by pathologists using tissue biopsy images are central for many tasks such as the detection of cancer and estimation of its current stage [2]. Known as histopathology, it is defined as the study of disease using microscopic examination of a specimen processed with methods like dehydration, sectioning, staining and fixed to a glass slide [6]. Recently, this analysis has become digital with Whole Slide Imaging (WSI) allowing users to capture entire slides as colored high resolution images [6]. It is important to note that color is especially important in histopathology, since different stains are used to highlight cell structures. The intensities of the stains give different information due to interactions with the cell components and metabolic processes.

One routine yet important step within histological analyses is the segmentation of nuclei. Nuclear morphology is an important indicator of the grade of cancer and the stage of its progression [3]. It has also been shown to be a predictor of cancer outcome [4] and counting mitosis events in nuclei is a useful marker of cancerous growth. [2] Currently, histological analysis such as these are done manually, with pathologists counting and evaluating cells by inspection. Developing automated methods to perform this analysis will help pathologists maintain consistent quality, allow for greater use of histological analysis by reducing cost and throughput - particularly in longitudinal studies, and allow for integration of results in electronic decision support systems.

However, automating nuclei detection is not a trivial task and can be challenging for a number of reasons. To fully understand the challenges, it is useful to detail the image creation process. Tissue biopsies are taken from an area of interest. The tissue is dehydrated, sectioned, stained, and photographed under a high magnification microscope (20X-40X). A light source illuminates the bottom of the sample and a sensor reads the relative absorption of the light through the sample. Cell samples are typically clear, that is, they allow light to pass through unaltered so stains are applied to the samples to allow for better differentiation of key cellular features. Eosin and hematoxylin are commonly used stains with the first staining nucleic acids a purple-blue and the latter staining proteins pink.

There are sources of error at each stage in the process. The tissue sample itself may have clumped nuclei or dividing those that overlap. Stain manufacturing and aging can lead to differences in applied color. It could also be the result of variation in tissue preparation (dye concentration, evenness of the cut, presence of foreign artifacts or damage to the tissue sample), stain reactivity or image acquisition (image compression artifacts, presence of digital noise, specific features of the slide scanner). Each stain has different absorption characteristics(sometimes overlapping) which impact the resulting slide color. Finally, storage of the slide samples can have aging effects that alter the color content. However, unlike medical fields like radiology, digital approaches to histological evaluation are fairly new so there isn’t always the same level of maturity in comparing scans of different origin. [2] [5]

Radiologists have established standards (such as DICOM) to ensure consistency between scans from different origins and time. Most relevant to this project, DICOM data files, contain metadata that allow users to easily manipulate raw image values to standard units based on specific machine characteristics. Ideally, histopathology would also work within a framework like DICOM where images can be standardized against experimental condition to ensure consistency across datasets.

The impact of such a standardization would be great. Aside from clinical and research applications, standardizing image analysis solutions to handle histological data from a variety of sources would also be valuable to the pharmaceutical industry. Recently, there has been considerable interest in the application of novel machine learning tools such as deep learning to aid in segmentation. Some have used grayscale images to perform automated slide analysis, but this ignores a deep source of data available from the stains. Color-based models generally work on raw pixel values, but could achieve greater accuracy through reducing the variance contributed by slide and stain specific variables. However, the approach must not be too general or else false positives will occur through reduction or altering the image signal. [1] [3]

## 2 Methods

The aim of this project is to address the impact of variability in histological images on the accuracy of segmentation algorithms. Image statistics were found on the raw images, and a CNN for nuclei segmentation was trained and tested to get a baseline. Four stain normalization techniques, histogram equalization, Reinhard, Macenko, and Kahn were then applied as means to reduce color variability of the raw images. The CNN was trained and tested again to get a final segmentation accuracy. This paper is unique in that it employs a wide variety of normalization methods, uses nuclei segmentation accuracy a metric, and tests the model on a different dataset to understand model generalizability.

**Fig. 1.**
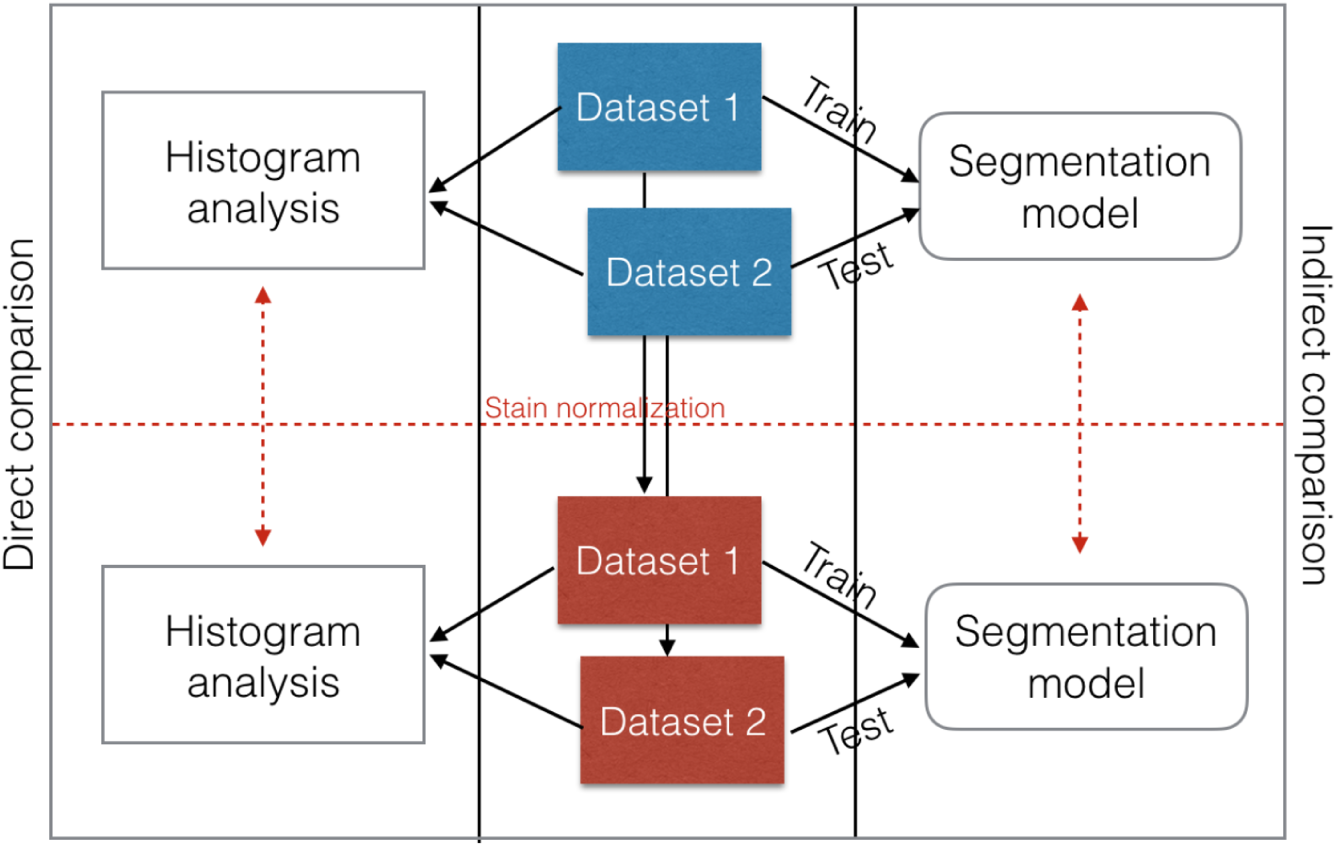
This is a schematic overview of our overall approach. We first used a direct histogram analysis to analyze the impact of stain normalization. We then trained a deep learning model on one dataset and tested it on another. We performed segmentation both before and after stain normalization to assess impact

## 3 Dataset

Multiple publicly available datasets were used in this analysis. 58 histological breast images were taken from the UCSB Bio-segmentation benchmark dataset. Each RGB image is 896x768 pixels was H&E stained and included a corresponding malignant or benign label. Manually annotated segmentation masks are also provided as a ground truth. 143 breast images were also used from the Case Western Reserve digital pathology dataset. Each RGB image was 2000x2000 pixels and was H&E stained. Manually annotated masks were provided for over 12000 nuclei.

These datasets were chosen because there were large differences in the resolution, color profile, and slide preparation quality. Obtaining such different images highlighted the problem facing automation, but also offered an opportunity to evaluate whether the segmentation classifier is generalizable outside of its training and validation set.

## 4 Stain Color Normalization

Stain normalization techniques involve transforming image pixel values. There are a wide array of techniques in literature, but most involve statistical transformations of images in various color spaces. This paper provides an overview of the four techniques used.

University of Warwick’s normalization Matlab toolbox contained example implementations of many of these techniques and served as the foundation for processing the training and test datasets. A training image was selected for training and test processing by inspection.

### 4.1 Histogram equalization

Histogram equalization is a commonly used image processing technique that transforms one histogram by spreading out its distribution to increase image contrast. In this analysis, histogram equalization was performed on each RGB channel in Matlab, effectively normalizing the color intensities frequencies between two images. However, histogram equalization is known to increase the contrast of noise and introduce artifacts since the assumption that the proportion of stains is constant between the images[kahn].

### 4.2 Macenko color normalization

The Macenko color normalization method transforms the images to a stain color space by estimating the stain vectors then normalizes the stain intensities. Quantifying the stain color vectors (the RGB contribution of each stain) provides a more robust means of manipulating color information. However, due to the variability of stains discussed previously, an automated method to determine the stain vectors is advised.

Macenko implements a method for automatic stain vector creation by considering images in the OD (optical density) space. Resulting OD values are a linear combination of the stain vectors. Low OD values are also thresholded to improve performance since background is considered to have an OD of 0. The largest OD values are found via a single value decomposition (SVD). A plane is calculated from the largest maximum values found, and the remainder of OD pixels are projected onto this plane with their vector lengths normalized to unit length. The angle of each point with respect to the first stain vector is calculated. To account for noise, the 1st and 99th percentile of pixels for each value are used to ensure a valid extremes. The extremes are then converted back to OD space. Intensity variation is corrected for each stain. An intensity histogram is plotted and the 99th percentile used as the maximum to account for noise. The target image stain intensity histograms are then scaled to the source maximum.

This method has been criticized for altering fully saturated and empty OD values during transformation and for altering the OD of both source and target images[kahn].

### 4.3 Reinhard color normalization

Reinhard color normalization aims to make one image ‘feel’ like another by transforming one image distribution to be closer to another. Reinhard transforms the target and source images into l” color space. In RGB color space, color channels typically have a high degree of correlation (ie. if there is blue there is likely green). Thus, any modifications to pixel values should have some impact on the other channels. L” color space was created to minimize the correlation between channels for natural scenes. L is an achromatic channel and while” are opposing Yellow-Blue and Red-Green channels.

After the source and target images are in L” colorspace, descriptive statistics are used to transform the target image’s’ colorspace. For each channel, the mean value is subtracted from the data points and the data points are scaled by the std of the target/std of the source. Finally, the average channel values of the source are added back to the data points. For images that are very different(a lot of sky vs a lot of grass), the above method does not produce accurate results. Reinhard suggests additional statistical methods to deal with widely varying images.

Instead of computing mean and standard deviation for on each channel for the entire image, the statistics are calculated for scene swatches or clusters (ie sky and grass). The distances are calculated to each cluster and divided by the standard deviation. The transformed pixels for each cluster are weighted by the normalized distances to produce a final output color. The technique easily scales if there are greater than two clusters. After the color transformation in L” color space, it is transformed back to RGB colorspace.

### 4.4 Kahn color normalization

Conceptually, the Kahn normalization technique is similar to the Macenko technique in that it estimates the stain vectors, deconvolve the image, maps the stain intensity to a target image, before reconstructing back in RGB colorspace. Kahn makes contributions in automatic stain vector calculation using a classifier with global and local pixel value information, and a non-linear stain normalization method.

The stain classifier uses SCD to split up image into color batches using oct-tree quantization. The SCD has its dimensionality reduced via principal component analysis as a color description of the full image, encompassing information on the stains. A relevance vector machine (RVM) supervised classifier had the best performance. It was also found that without the global SCD information, the classifier assigned significant probability to only weakly stained classes.

Optical density values are analyzed for each channel after being separated by the classifier (stains, and background). Statistics were calculated for the stain channel distributions with OD outliers excluded. A spline function was then created to map the channel statistics of one image to another. The spline ensures was created to ensure that the maximum and minimum OD remain unchanged through the mapping. After the transformation the image is converted by the RGB colorspace.

**Fig. 2.**
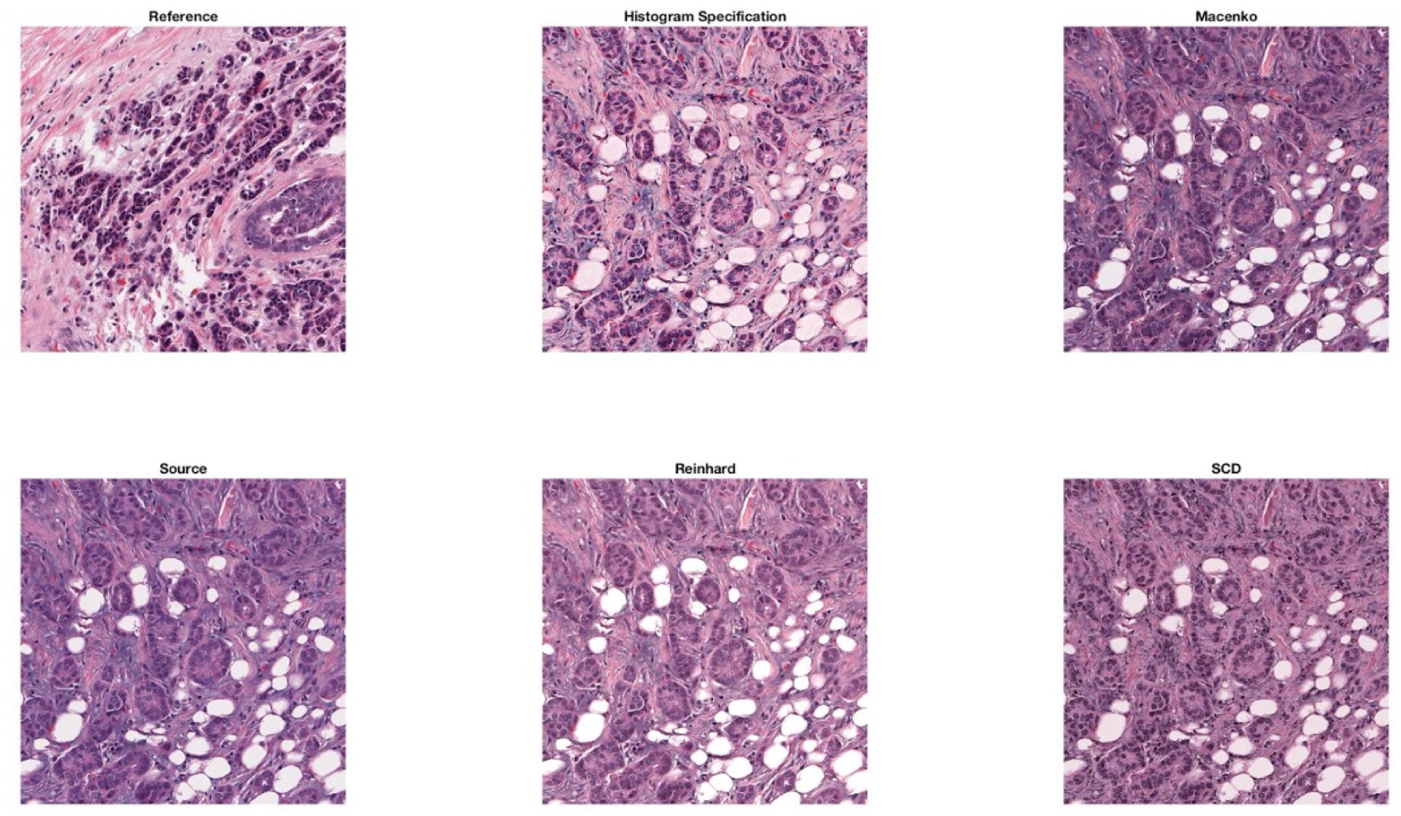
Comparison of stain normalization techniques on a sample training image. Source image was normalized to reference image using histogram equalization, Reinhard, Macenko, and Kahn methods respectively

## 5 Stain Normalization

Stain normalization techniques involve transforming image pixel values. There are a wide array of techniques in literature, but most involve statistical transformations of images in various color spaces. This paper provides an overview of the four techniques used. University of Warwick’s normalization Matlab toolbox contained example implementations of many of these techniques and served as the foundation for processing the training and test datasets.

**Fig. 3.**
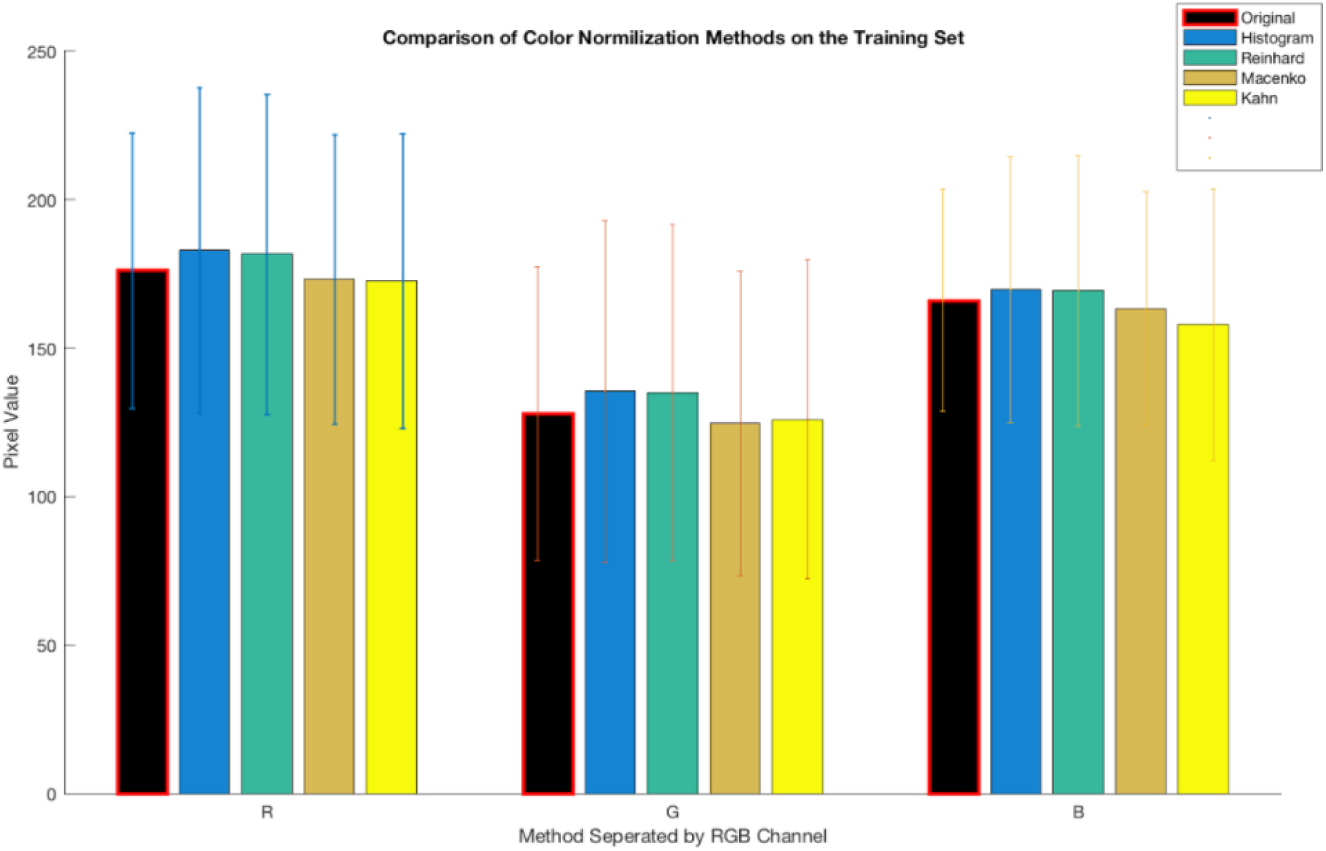
Comparison of stain normalization techniques on Case Western Reserve training set. Mean pixel value and standard deviation was calculated for each color channel demonstrating a quantitative change in color composition and increased variance.

## 6 Image segmentation using CNNs

This section describes the methods used to generate classifier models. The goal was to train a model using one dataset and to test using another. We aimed to perform this approach using different normalization strategies to narrow down on an approach that best reduced variability and improved performance

### 6.1 Model selection

We first trained and validated the model on the same dataset to make sure that our training procedure was working correctly. For this we used breast tissue slices from [3]. We split the dataset into 70% train and 30% validation. We then tried a number of different network architectures, dataset augmentation techniques and patch selection strategies to narrow down on a working approach. These efforts have been briefly outlined in Table X. We chose to use a fully convolutional network instead of a regular fully connected network to enhance throughput and quickly retune the network based on results. We used the Caffe deep learning framework to design these models. The final architecture that was chosen after the validation procedure for this problem was (Conv-BNorm-ReLU)x6 - (Full ConV) - Softmax [3]. It has extra convolutional layers as compared to a regular Alexnet but only one ‘fully connected’ layer (that has in this case been transformed into a fully convolutional layer).

### 6.2 Training Dataset

The training dataset consisted of histological sections of breast tissue. These were all standardized at a resolution of 2000x2000 and a magnification of 20x. We sampled patches from these images in order to develop a pixel level classifier. Patch sizes of 32x32 and 64x64 were experimented with, and we found that a patch size of 64x64 worked best for training. Some of the models experimented with have been listed in Table X.

However, on training we realized that randomly sampling from non-nuclear regions as defined by the hand annotations was not sufficient. This is because the annotation mask only accounted for a subset of all the nuclei in the image. Thus, there was a significant probability of sampling unannotated nuclei while developing patches for the training set. To address this problem, we used the approach outlined by [3]. Nuclei are known to absorb greater levels of the eosin (red) stain and so the red channel in the images was enhanced. A negative mask was thus generated defining regions outside of these enhanced red zones that were deemed safe for non-nuclei class patch selection. We also made sure to allocated a third of the non-nuclei patches to boundaries around the nuclei. This ensured that the algorithm was careful to clearly mark out these boundaries and not club nuclei together. The model prediction accuracy benefitted from both these approaches.

The test set was composed also of breast tissue slices from a dataset provided by the BioImaging lab at UCSB [link]. These were of a much smaller resolution (896 x 768) and are referred to as the the BioSegmentation benchmark. This dataset proved to be exceptionally useful for this problem because the images were quite different from our training set both in terms of image quality and resolution and also in terms of the staining used (more eosin content). This was ideal for our approach since we could see how well the features learnt on a moderately different dataset could transfer to this problem, and most importantly how we could augment that process through better normalization and standardization.

Patching was not required for the test set because we were using a fully convolutional network. This meant that our model could simply process an entire image in one pass instead of needing it to be broken down into pixelwise patches which involves a lot of redundant and time consuming computation. We used the approach first described by [15].

### 6.3 Training

Once our model architecture and dataset generation approach had been finalized, we began to train separate model for each of the normalization scenarios as shown in Fig 4. We used a batch size of 1000 because that could fit comfortably in our memory (P100 GPU 16GB x 2) and also leave enough room for sharing with other runs. Any further increases in batch size would not result in a performance increase because of memory bandwidth limitations. We initially tested architectures with a considerably large dataset of 2,500,000 images. However, we soon realized that there was a critical flaw in how the binary LMDB databases were being generated which was corrupting all those values, thus we chose to use a slower I/O procedure and a smaller dataset. Since the images from which patching was performed were still the same, we did not feel that the results would be adversely affected by this. From previous experience, the authors expect that merely oversampling from the same dataset would have improved accuracy by not more than a few percentage points, whereas more structured sampling like generating a negative annotation mask as described previously can have a more significant impact.

**Fig. 4.**
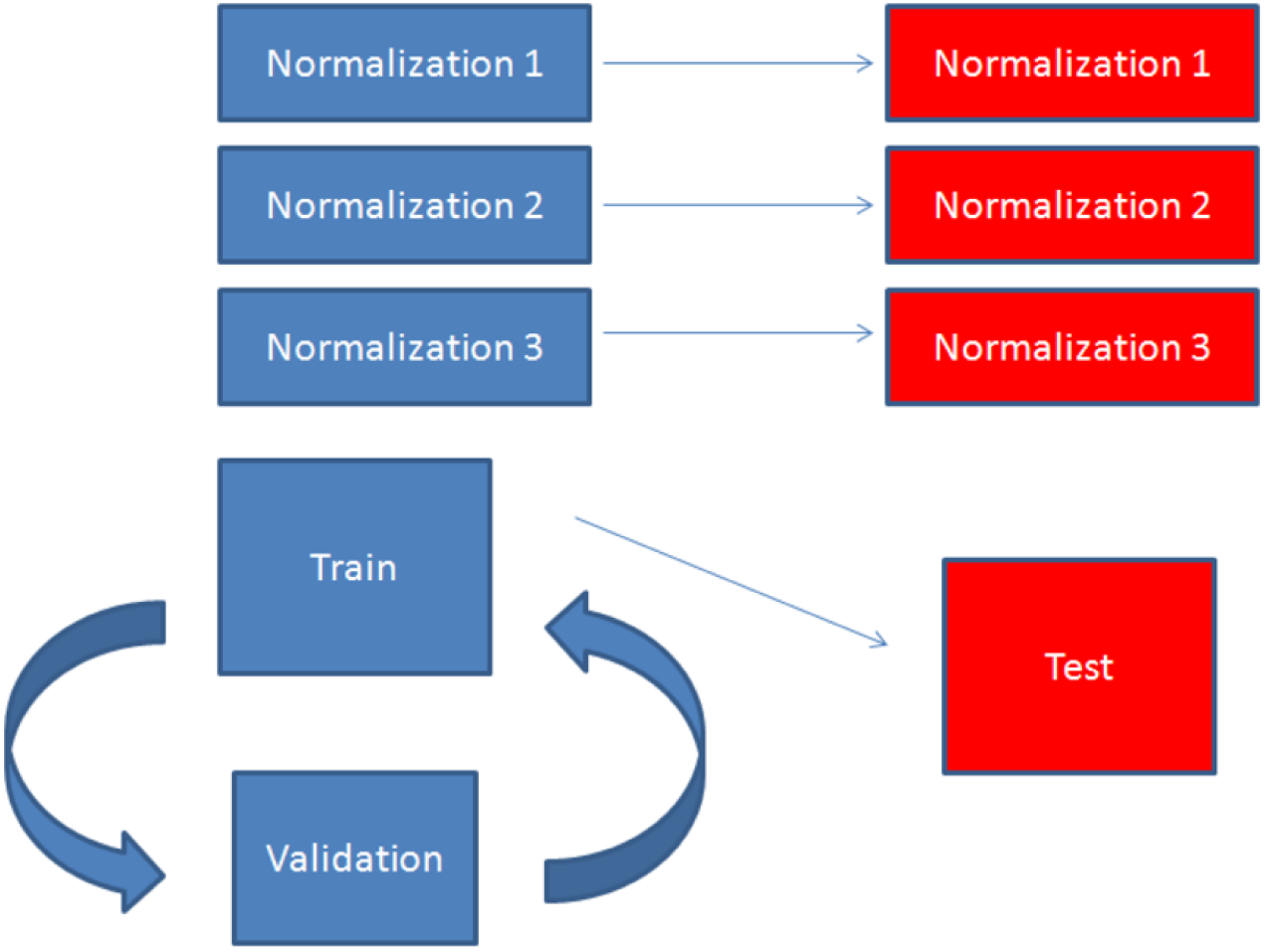
Outline of model validation and testing procedure

**Fig. 5.**
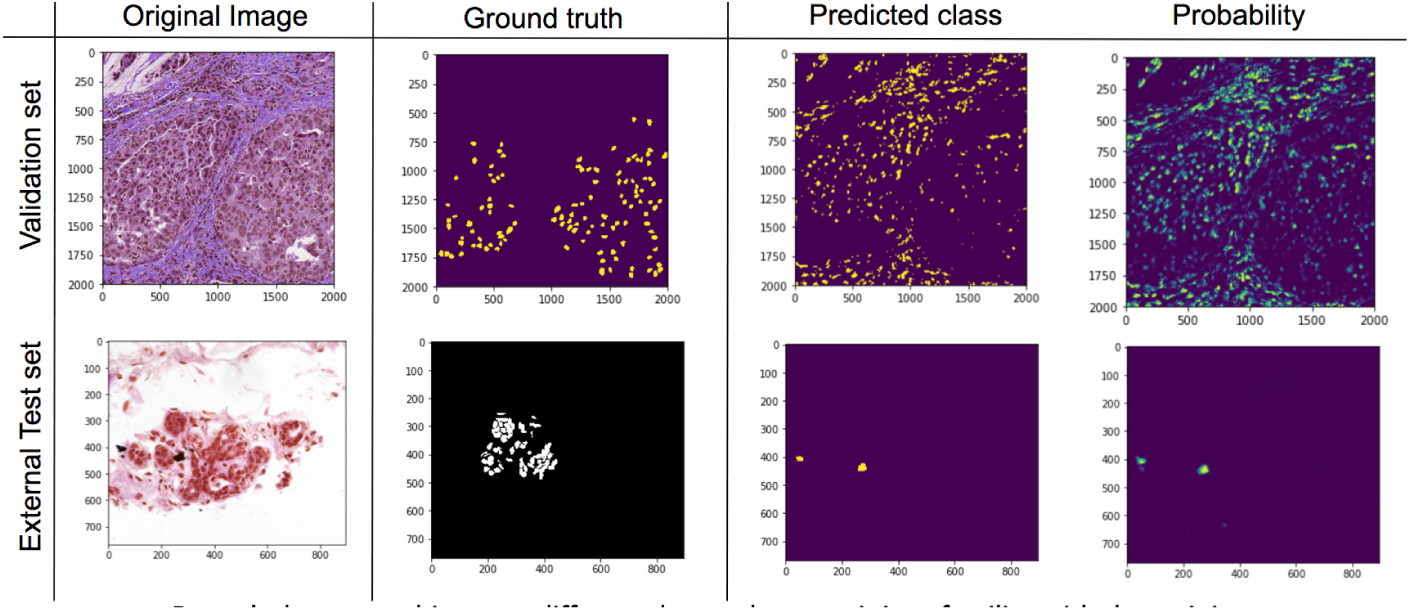
Comparison of normalized and unnormalized models

Most of the models would quickly converge within 5-10 epochs. We allowed each normalization model to run up to 25 epochs just to be sure. There were four models - these corresponded to the four techniques outlined previously: Histogram Equalization (SH), Macenko (MM), Reinhard (RH), Spline Mapping (SM). There was a also of course a model for the unnormalized (Unnorm) case

## 7 Results

Results from the segmentation model were analyzed both visually and using quantitative metrics.

### 7.1 Visual inspection

The top row in Fig 6 shows the original images after being transformed using the four different stain normalization approaches. We can see that all four images appear different in some respect. For example, HE and RH, which involve stain normalization through working directly with the color values show a noticeable blue tint. This is more pronounced in HE, where non-nuclear regions in the top right of the cell get quite heavily stained with hematoxylin. On the other hand, SM and RH, which both use stain vectors to map an input image to a target space, don’t show a blue-ish tint and provide a much more robust transformation that is true to the semantic information of the parent image (eg: treating nuclei and background regions distinctly)

**Fig. 6.**
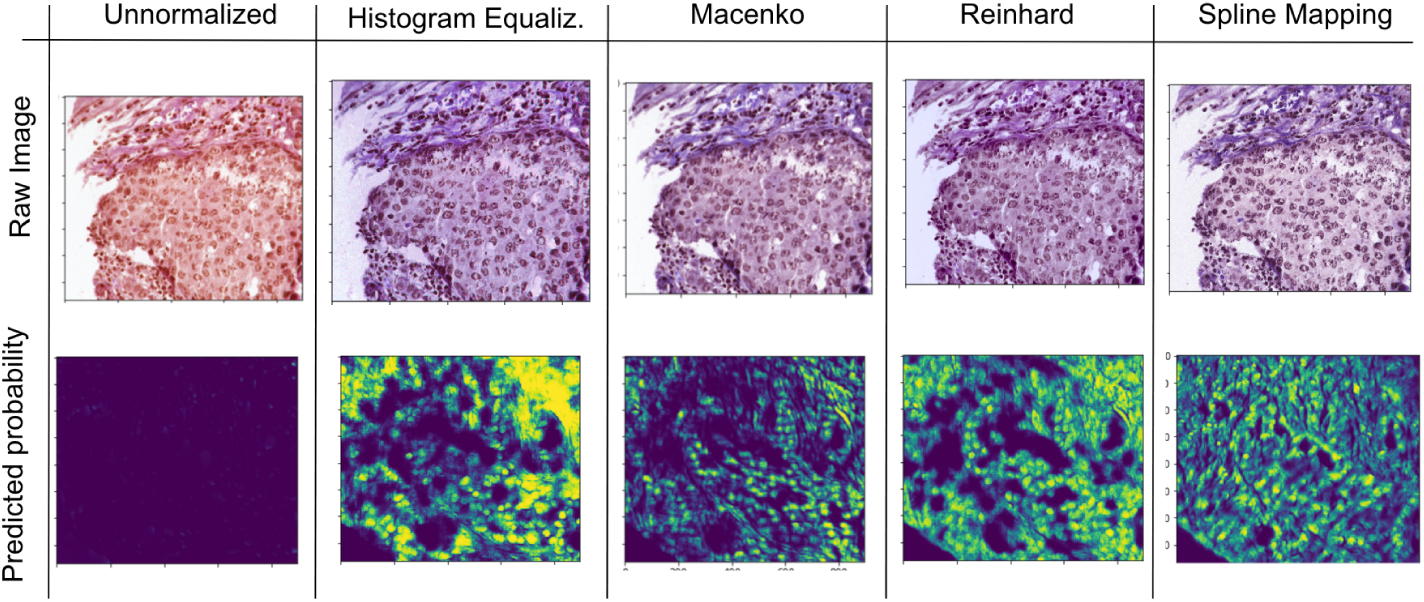
Comparison of different stain normalization procedures on model performance on test set

The bottom row looks at the class probability for nuclear regions as predicted by models trained on datasets that were each stain normalized differently. Clearly all four normalized sets perform far better than the unnormalized dataset where almost no nuclei were detected due to the drastic change in staining as compared to what the model had been trained on. HE does pick up most of the nuclei but also a lot of background noise due to its inability to differentiate clearly between different types of staining. RH is also more sensitive to noise but does a better and clearer detection of nuclei as is visible in the clear boundaries. SM clearly performs the best at segmenting nuclei while also being most robust to false positives.

### 7.2 Quantitative assessment

To perform a more rigorous quantitative assessment, we looked at a range of different metrics calculated over a randomly selected set of 15 test images. This was essential because simply calculating classification accuracy would be insufficient for this sort of segmentation problem. For instance, even if a classifier were to only classify pixels as non-nuclear regions it would still be around 85-90% accurate because the vast majority of pixels don’t lie within nuclei. Thus, the range of metrics that we calculated helped to characterize model performance from many different perspectives.

Given the set of all nuclear pixels in the set, recall tells us what fraction of those were picked up by the model. Clearly SM does a great job in this area. SH and RH also do well but when we look at their precision values they are not as high as those for SM. Precision measures how many of the positives that you picked up were actually relevant. This indicates the tendency of SH and RH to pick up more false positives than SM. This trade-off between true and false positives is best captured by the ROC curve (Fig 7). Here, we see that the unnormalized case doesn’t add any value at all while all the normalization scenarios show improved prediction accuracies. SM is the clear winner showing an excellent range of operation at a TPR of ¿80/90% while only allowing an FPR of 50%. This is very impressive considering how the model was trained on a staining visually very different from the one in the test data. This difference is quantiatively captured by the AUC. Finally, the F-score is another attempt to capture segmentation accuracy without getting bogged down by all the true negatives. It calculates the intersection of pixels that have been classified as nuclei in both the prediction and the ground truth and it divides that over the union of all pixels classified as nuclei by either set. Again, SM is seen to be the best at improving accuracy of the algorithm.

**Fig. 7.**
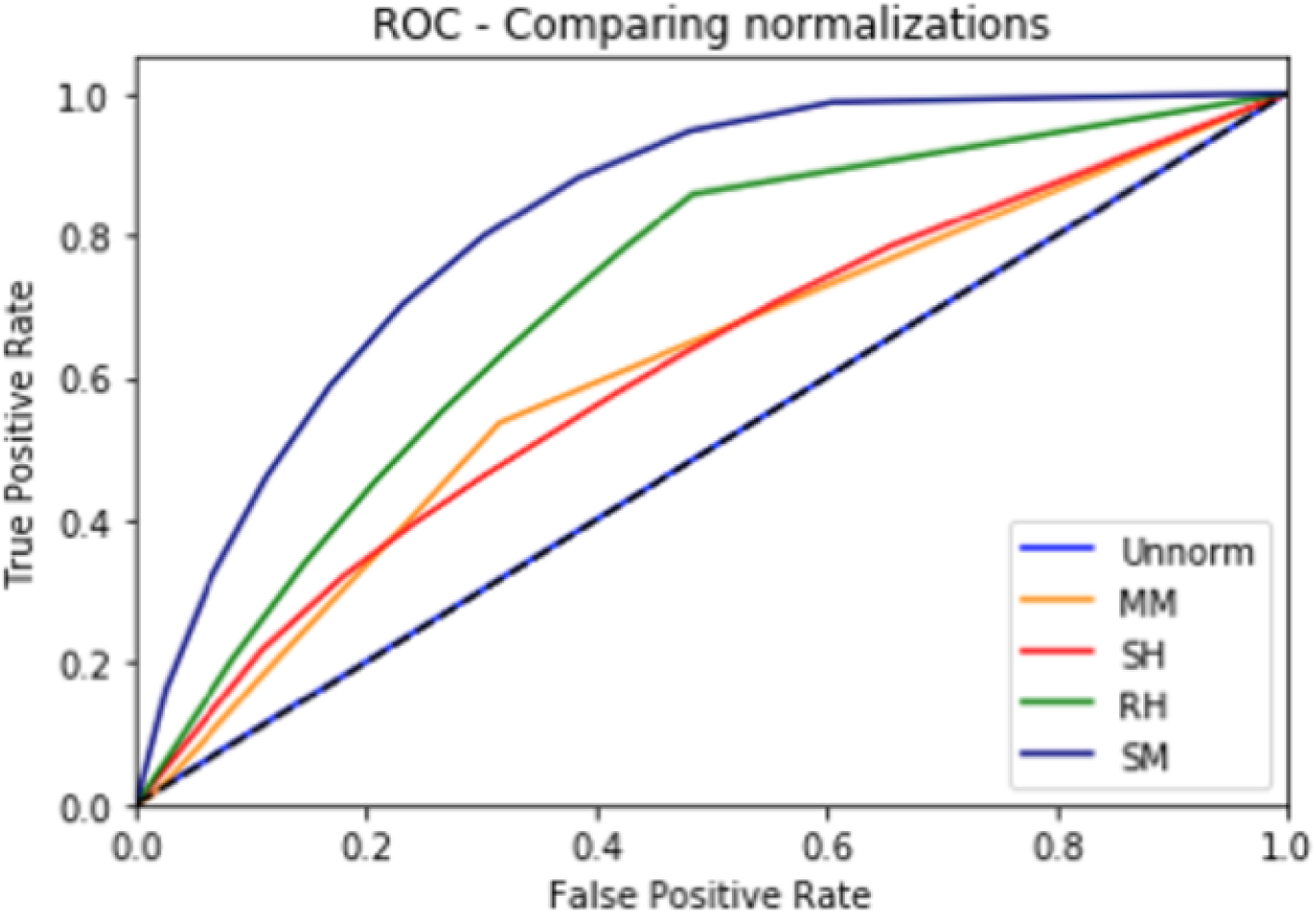
ROC Curve for models trained using different stain normalization schemes

While we aimed to run the same number of epochs for the same images for all the (4+1) normalization scenario models that we were testing, due to time constraints before the presentation we ran a few models for lesser than 25 epochs. As we attempted to update the accuracy numbers with newer figures, it led to an interesting finding that the accuracy tended to decrease with increasing number of epochs. This is quite possible given that increased number of epochs leads to a greater chance of overfitting to the training dataset. The goal of our approach is to develop a generalizable approach to move across datasets and so it would be more sensitive to overfitting than a regular model that works on similar looking images.

**Fig. 8.**
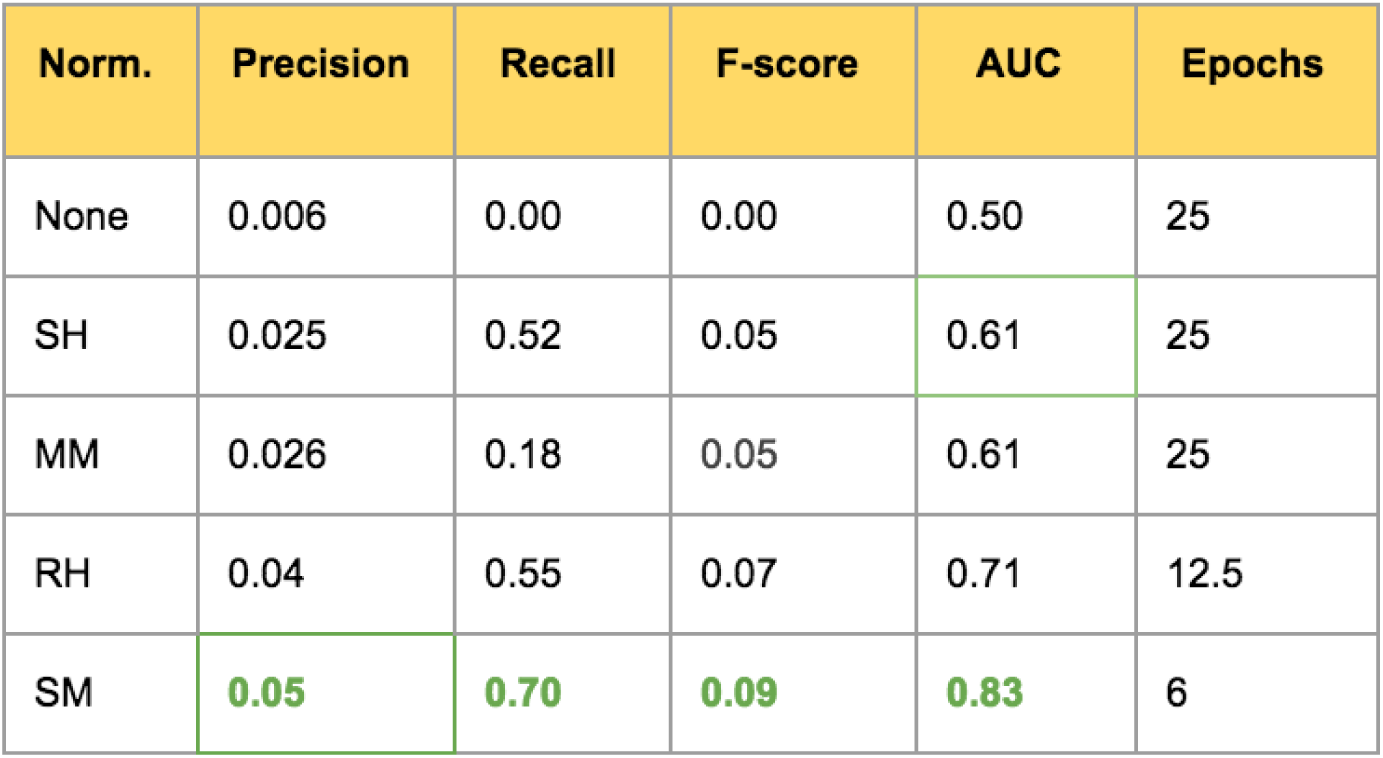
Quantitative comparison of model performance where training data has undergone different stain normalization procedures

## 8 Discussion

Through this study, we have explored several stain normalization approaches that were all shown to reduce inter slide variability. The results (particularly AUC, F-score) clearly indicate that using a stain normalization approach increases the performance of the deep learning based segmentation algorithm. We found that SM performed better than all other approaches. We believe this is because it use a non-linear mapping function that is more accurate than the other approaches. It is able to delineate between different regions and map them appropriately to the target space.

We also noticed that the model seems to perform more poorly in case of normalizations that tend to be biased more towards the eosin channel. In future, it may make sense to normalize the stain of the training dataset using two different approaches. This would push the model to become robust to these subtle changes and be less dependent on any one channel. In fact, this could also be looked at as a regularization approach to enhance generalizability of deep learning based models in this space and prevent overfitting. On the other hand, we must remain conscious of the fact that staining color is a very valuable source of information in histological analyses and reducing the model’s recognition of critical differences would prove detrimental. Thus, there is a balance to be achieved here.

## 9 Conclusion

In this study, we looked at the impact of stain normalization as a means of reducing inter-dataset variability in histological sections and also as a way to improve the accuracy of segmentation algorithms across datasets. We were able to quantify improvements in both domains. To the best of our knowledge, this is the first study that looks at all four of these techniques and assesses their usability in the context of deep learning based segmentation models.

